# Range expansion of the invasive barnacle *Balanus glandula* into the Wadden Sea, its habitat and parasite fauna compared to established barnacle species

**DOI:** 10.64898/2026.08.25.747026

**Authors:** Rosalinde van Ooijen, Rebecca Büring, Annika Cornelius, Haiyan He, Dick van Oevelen, David W. Thieltges, Cyril Hammoud

**Author notes:** Corresponding author David W. Thieltges.

## Abstract

The impact of invasive species on marine ecosystems is rapidly increasing, where they often outcompete native species in the absence of natural enemies. The parasite release hypothesis states that the success of invasive species relates partly to the loss of natural parasites during introduction and lower susceptibility to native parasites. Barnacles are highly successful invaders due to broad environmental tolerance and dispersal via shipping, but whether parasite release also participates in this success remains unknown. In this study, we analyse parasite infection patterns in native and invasive barnacles in the Wadden Sea by surveying communities across tidal zones. Additionally, year-round molecular monitoring of larval stages and a literature review were used to track the distribution of the invasive Pacific barnacle *Balanus glandula* in Europe and document its appearance in the Wadden Sea. The long-established invasive *Austrominius modestus* dominated the high and middle intertidal zone, whereas native species (*Balanus crenatus* and *Amphibalanus improvisus*) prevailed in lower zones. Native and invasive barnacles differed in parasite infection frequency (mostly cestodes and trematodes). The native *Semibalanus balanoides* had the highest prevalence (27%), followed by the invasive *A. modestus* (11%), and no infections were found in *B. glandula*. Lower parasite prevalence in invasive barnacles is consistent with the hypothesis that parasite release supports invasion success. In the absence of competent parasites, *B. glandula* could impact native barnacles through competition. Continued monitoring of *B. glandula* is recommended to track its distribution, interactions with native species, and parasite acquisition, providing further insight into the parasite release hypothesis.

## Introduction

In recent decades, anthropogenic activities have accelerated the pace of biological invasions in marine ecosystems globally (Occhipinti-Ambrogi and Savini 2003). Shipping activities have increased drastically, both in the amount of shipping and in the distances travelled, allowing species to be introduced to new ecosystems on a global scale (Seebens et al. 2013). Species growing on hard substrates can be transported by attaching to the hull of ships, a phenomenon called hull fouling (Sylvester et al. 2011; Clarke Murray et al. 2012). Marine invertebrates that have a larval stage, called meroplankton, such as molluscs, crustaceans, echinoderms, and cnidarians, can also be transported in the ballast water of ships, with over 12 billion tons of ballast water being discharged yearly upon arrival at the destination port (Bax et al. 2003). Aquaculture is another important contributor to the introduction of species into new ecosystems. Species intentionally introduced solely for aquaculture practices can escape from farm settings and establish populations in the wild. In addition, other species can be co-introduced unintentionally with aquaculture transfers (Oficialdegui et al. 2025). Although antifouling paints and ballast water treatment in shipping, as well as biosecurity measures in aquaculture, increasingly help reduce introductions, there is still an increase in the numbers of species introductions in marine ecosystems (Seebens et al. 2013).

Once introduced in a new ecosystem, species need to first develop self-sustaining populations to become established. A lack of natural enemies, such as predators and parasites, may often facilitate establishment, known as the enemy release hypothesis (Torchin et al. 2001; Liu and Stiling 2006; Dunn et al. 2023). The parasite release hypothesis specifically proposes that the loss or reduction of parasites during introduction can facilitate the establishment and spread of invasive hosts (Torchin et al. 2002; Goedknegt et al. 2016). In many cases, established species can also spread from their original point of introduction and lead to secondary introductions in other areas, thus becoming invasive and extending their range in recipient ecosystems (Blackburn et al. 2011; Johnson et al. 2012). In these recipient ecosystems, spreading invasive species may have a diversity of impacts on resident biota (Vila et al. 2011). Some invasive species directly compete for space or resources with native species. Others, such as predators or grazers, can change the abundance of native prey and might cause trophic cascades, changing the overall community structure (Sammarco et al. 2015; Doherty et al. 2016; Brown et al. 2023), thereby affecting energy pathways in food webs (Baird et al. 2012). Finally, invaders may also affect parasite-host interactions in resident biota by introducing new parasites or diseases that can spread to native species, or by being susceptible to native parasites that can then spill back to native host populations (Goedknegt et al. 2016). As a consequence of these biological impacts, invasive species can degrade the ecological and economic values of ecosystems and are thus often considered a threat to marine biodiversity (Molnar et al. 2008; Geller et al. 2010).

Barnacles are among the most successful invaders in marine ecosystems and have been introduced to new regions all over the world, mainly aided by ship movement (Torres et al. 2012; Aguirre-Tellez et al. 2025). In Europe, the barnacle *Austrominius modestus* is a good example of a successful invader (Gallagher et al. 2016). This species is native to New Zealand, Australia, and Tasmania and was introduced in Europe in the 1940s, where it most likely arrived with ships during the Second World War (Allen et al. 2006). In some places, such as in the Wadden Sea, *A. modestus* has become more abundant than native barnacles such as *Semibalanus balanoides* and *Balanus crenatus* (Witte et al. 2010; Buschbaum et al. 2012; Reise et al. 2025). Although *A. modestus* was already introduced in the Wadden Sea in the 1950s, its population only dramatically increased in abundance after the climate became more suitable, with milder winters and warmer summers (Gittenberger et al. 2009; Witte et al. 2010). A more recent introduction is the barnacle *Balanus glandula*, which was first recorded in Europe on the coast of Belgium in 2015 (Kerckhof et al. 2018). *Balanus glandula* is native to the Pacific coast of North America but has a wide distribution as an introduced species to the coasts of Argentina, Japan and South Africa (Schwindt 2007; Geller et al. 2008; Laird and Griffiths 2008). The species has so far not been observed in the Wadden Sea, but given its geographical proximity to the area of first introduction and the species’ high dispersal potential via planktonic larvae, establishment in the Wadden Sea appears likely. *Balanus glandula* is believed to occupy the same niche as *Semibalanus balanoides* (Barnes and Barnes 1956). Indeed, as both species also occupy the same tidal zones and substrates, they will likely compete for space when co-occurring. Beyond uncertainties regarding the presence and distribution of *Balanus glandula* in the Wadden Sea, little is known about parasite infection patterns in native and invasive barnacles. Consequently, it remains unclear whether invasive barnacles experience reduced parasite pressure relative to native species, as predicted by the parasite release hypothesis. Addressing this knowledge gap is essential for understanding whether parasite release contributes to the establishment and spread of invasive barnacles in the Wadden Sea.

This study aimed to investigate the distribution of *Balanus glandula* in the Wadden Sea and to compare its habitat and parasite fauna with other resident (native and invasive) barnacle species. For this, we sampled barnacles at various locations in both the Dutch and the German Wadden Sea, complemented by metabarcoding of larval stages in the Dutch Wadden Sea. In addition, we investigated the current distribution of *B. glandula* in Europe, based on information from other sampling programmes and records from the scientific literature.

## Material and methods

### Study area

The Wadden Sea, designated as a UNESCO World Heritage Site, is one of the world’s largest and most dynamic intertidal ecosystems, characterised by unique geomorphological and ecological processes and spanning 500 kilometres along the Netherlands, Germany and Denmark. The Wadden Sea has many microhabitats due to its environmental gradients and different tidal zones, which create habitats for species that are specialised for extreme conditions (Reise et al. 2010; Baptist et al. 2019). This variety of habitats supports complex food webs and high biodiversity, offering a natural setting to explore ecosystem functioning. The Wadden Sea also provides important ecosystem services, including carbon storage, coastal protection, and breeding grounds for migratory birds, all of which are increasingly affected by human activities (Baptist et al. 2019).

### Field sampling in the Wadden Sea

The presence of mature *B. glandula* was studied by sampling barnacles from February 2025 till October 2025 at 12 locations along the Dutch and German coast of the Wadden Sea, including the coasts of the islands Texel and Sylt, and an artificial reef near the island Vlieland (53.00° – 55.04° N, 4.72° - 8.44° E) (Suppl. Fig S1). Samples were taken from various artificial and natural substrates. Artificial substrates included buoys and man-made structures such as concrete, steel from coastal protection structures and harbour walls. Barnacles on natural substrates were sampled in bays and along sandy coasts, and included the blue mussel *Mytilus edulis*, the Pacific oyster *Crassostrea gigas*, stones and wood.

A total of barnacles 928 was collected across all locations (range 4 to 143 barnacles per site) and taken to the laboratory for species determination based on morphological characteristics, if possible, still attached to the substrate (Suppl. Table S1). Otherwise, barnacles were carefully scraped from the substrate using a surgical blade. Collected specimens were kept at 4 °C without water, if needed, for a few weeks, until determination. The shell diameter of each specimen was measured before proceeding with identification according to Hayward and Ryland (2017) and Kerckhof et al. (2018). Morphological characteristics used for species identification include the number of wall plates, the structure of the inside of the wall plates, the presence of a calcareous basal plate, the shape of the tergum and the scutum and the presence of a black band on the scutum.

After species determination, the 928 barnacles were dissected and screened for parasites. Specimens were removed from their calcareous shell plates from the back and laid out on a slide, using tweezers and a needle point. Barnacle tissue was screened for associated fauna with a Zeiss Discovery V8 stereomicroscope with a 1.5x FWD 28 mm lens. Associated fauna, defined here as organisms found in barnacle tissue independent of confirmed parasitic status, were identified at broad taxonomic level (gregarines, trematodes, cestodes, planarians and nematodes) based on Colston (2012) and Chuang (2021). The number of specimens was counted from each infected barnacle, and pictures of representative specimens were taken with a Zeiss Axioscope 5 stereomicroscope.

In addition, the relative abundance and habitat of barnacle species occurring in the Wadden Sea were studied in ten out of twelve sites. Due to time constraints, the remaining two sites (Mokbaai and Oudeschild) were not included in this component of the study. At five locations in the Netherlands and five in Germany, the sites were divided into ‘high intertidal’, ‘middle intertidal’ and ‘low intertidal’ or ‘subtidal’ as barnacle species occur in different tidal zones. Depending on the accessibility of the area and the tide, the different locations were sampled during low tide. The low intertidal zone was identified based on its position closest to the waterline during low tide. The middle and high intertidal zones were distinguished by their elevation relative to the low intertidal zone and their degree of exposure at the time of sampling. Barnacles from the subtidal could be sampled from washed-up substrates and from an artificial reef structure that was taken back to shore (Dickson et al. 2023). Pictures of randomly placed quadrats (10 x 10 cm) were taken at ten different sites and four tidal zones (n = 137). A subsample of barnacles was identified to species level after which the barnacles in the pictures were counted manually using the open-source image processing software ImageJ (Schneider et al. 2012). The relative abundance of barnacle species per tidal zone and per sample location was then calculated.

### Metabarcoding of barnacle larvae DNA

Barnacle larvae were monitored from November 2023 to October 2025 at the NIOZ jetty on Texel (53° 00 ‘06.5”N 4° 47’ 20.5”E). During each sampling event, two 50 L seawater samples were collected from the jetty using buckets. The collected seawater was first pre-sieved through a 3 mm mesh to remove larger organisms, such as jellyfish. The remaining water was then filtered through a 150 µm mesh net to retain barnacle larvae and other planktonic organisms. The resulting samples were preserved in DESS and stored until laboratory analysis at Wageningen Environmental Research. All DESS-preserved samples were processed for DNA extraction using the DNeasy PowerSoil Pro Kit (Qiagen). Metabarcoding libraries were prepared using the CO1 marker following a two-step PCR protocol and sequenced on an Illumina NovaSeq platform. Taxonomic assignment was performed using reference databases (BOLD and NCBI GenBank), after which sequence data were quality-filtered. All sequences with fewer than ten reads in total were removed from the data, and the maximum remaining read count of a sequence in the negative control was subtracted from all samples. To account for differences in sequencing depth, all samples were rarefied for each marker, resulting in a sequencing depth of 280k reads. CO1 reads were averaged per month per taxon, and zero reads were excluded from the visualisation (Büring et al., manuscript in preparation).

### European distribution of *B. glandula*

For the overall *B. glandula* distribution in Europe, a literature search was performed on the Web of Science Collection. The search was performed using the Topic field (title, abstract, and keywords), combining the species name with geographic keywords corresponding to European countries with a coastline. Records of *B. glandula* in Europe were also searched for in biodiversity databases such as the Global Biodiversity Information Facility but were not included in the final results due to excessive uncertainty in the identification of the specimens reported. An additional unpublished thesis focused on *B. glandula* distribution in Belgium (Bouwens 2019) was included in the overview. Latitude, longitude and location name were documented for the records found. In addition, results from the expanded Rapid Assessment Survey (eRAS) from 2025 were included. Settlement plates are deployed annually, and plate lines consisting of three plates per line remained in the water for 12 weeks. This method is supplemented by the Rapid Assessment. In late summer, various habitats (soft bottoms, floating docks, hard substrates) are examined for the presence of species (Hoppe et al. 2016). All records of *B. glandula* in Europe were plotted on a map, as well as known absence of *B. glandula* (Suppl. Table S2).

### Data analysis

To assess whether infection prevalence differed between native and invasive barnacle species, a binomial regression was fitted with infection status (infected vs. not infected) as the dependent variable and native status (native vs. non-native) as the predictor. A log-likelihood ratio test was used to compare the regression model to the null model, as this approach is considered more accurate than Wald’s method (Whitlock and Schluter 2020). To improve model performance and avoid issues associated with zero counts, a value of one infection was added to the group *B. glandula*, which initially had zero infections. A second binomial regression was fitted and evaluated in a similar way to examine the association between barnacle species as the predictor and infection status as the dependent variable. Post hoc pairwise comparisons were performed using Tukey-adjusted estimated marginal means to assess differences in infection probability among species.

## Results

### 1. Distribution of *Balanus glandula* in Europe and in the Wadden Sea

Field sampling on Texel and on the coast of the Dutch mainland provided the first record of *B. glandula* in the Wadden Sea (84 specimens in total). This is the first time *B glandula* has been observed at such high latitude in Europe, after its initial introduction in Belgium and Europe in 2015 (Fig. 1). We found no specimens from sites located on the coast of the German mainland or on the island of Sylt. Morphological species identification was confirmed by the author of the first *B. glandula* record in Europe (Kerckhof, pers. com.)

**Fig. 1.**
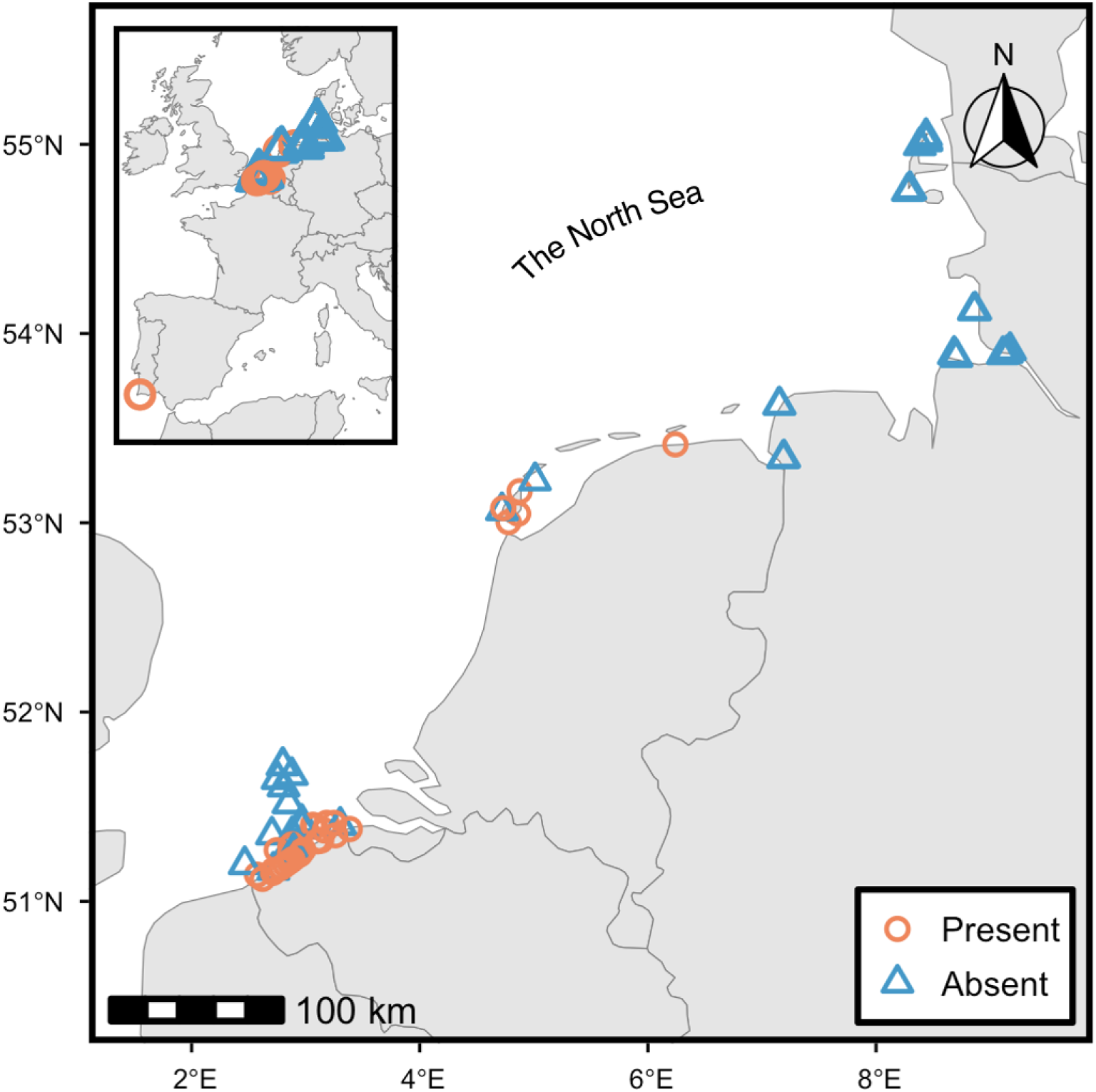
Distribution records of *Balanus glandula* and locations where the species was absent during investigations along the coast of Europe. Map data obtained from the Natural Earth dataset accessed via the rnaturalearth R package (Massicotte and South 2017).

At European scale, our literature search also identified what is likely another independent introduction of *B. glandula* in a European country, on an offshore mussel farm in Southern Portugal in 2018. No other records of *B. glandula* on the European coastline were found (Fig. 1).

### 2. Habitat preference of *Balanus glandula* and other barnacles in the Wadden Sea

*B. glandula* was present at several sampling locations in the Dutch Wadden Sea, including Mokbaai, the coast at Oudeschild and Cocksdorp, the coastal protection groyne at Paal 17 (all located on Texel), and on the dykes of Lauwersoog (located on the mainland) (Fig. 2). The species was characterised by lower relative abundance compared to most other barnacle species present at the sampling sites (Fig. 3b), and all individuals recorded were observed in the high intertidal zone (Fig. 3a).

**Fig. 2.**
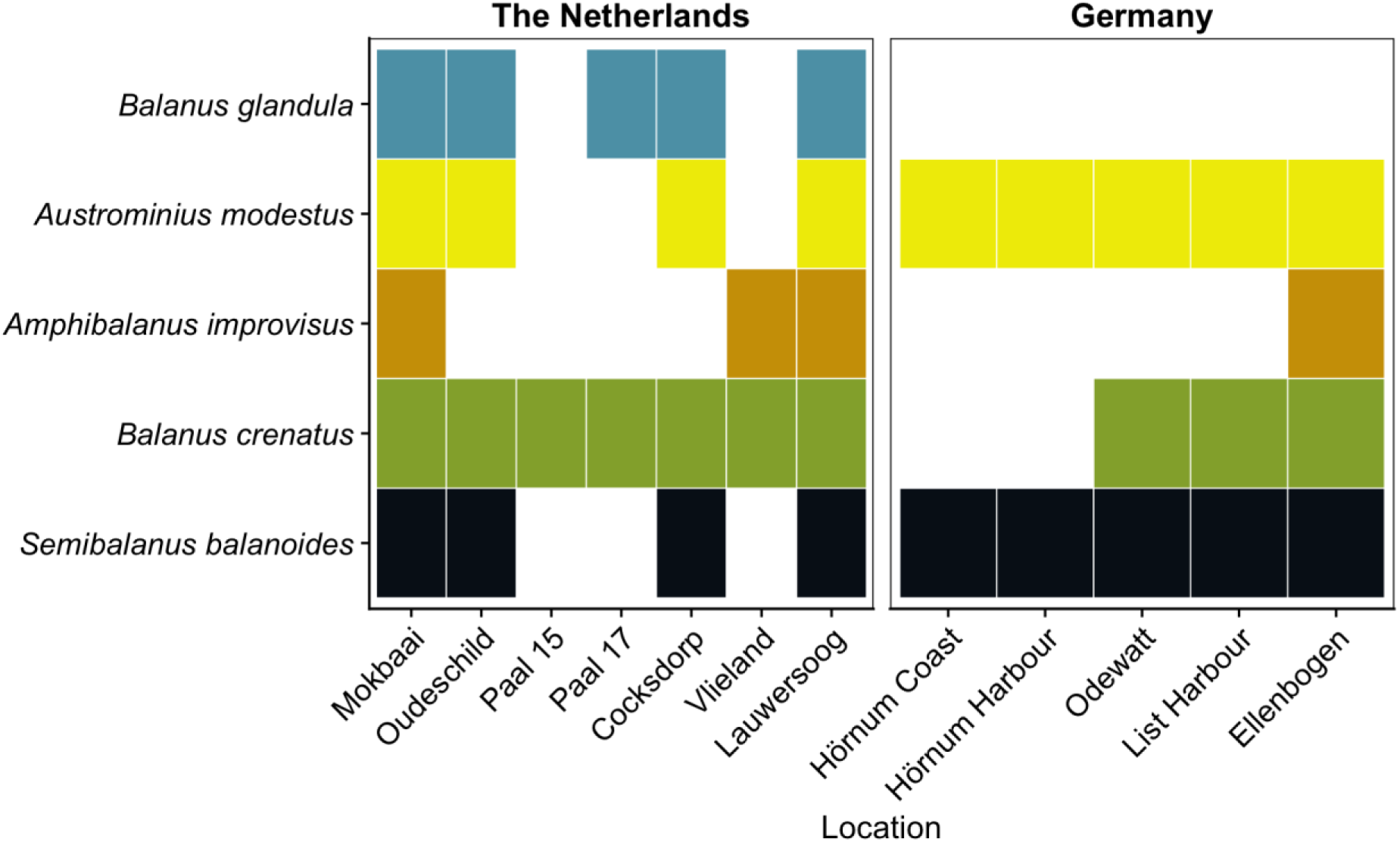
Distribution of *B. glandula* and other barnacle species along the Wadden Sea coast from South to North, based on the field sampling campaign in 2025.

**Fig. 3.**
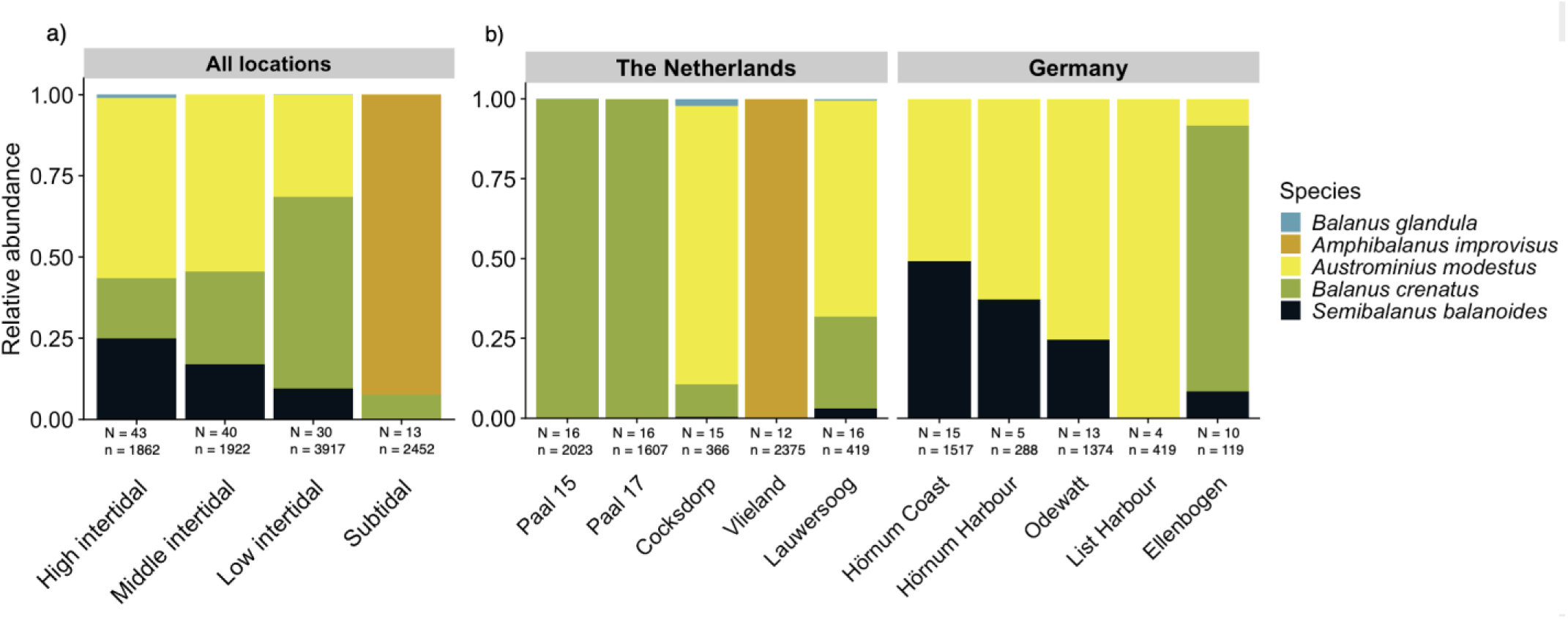
Relative abundance of barnacle species in the Wadden Sea a) at different tidal zones b) at different locations from south to north. Locations Mokbaai and Oudeschild were not sampled for abundance. The two panels are not based on identical datasets: panel a includes location ‘Oost’ and excludes ‘List Harbour’ and ‘Hornum Harbour’. N represents the number of quadrats taken, and n represents the total number of barnacles.

The high intertidal zone was mostly dominated by the invasive *A. modestus*, followed by the native *S. balanoides*. In the middle intertidal zone, *A. modestus* was also dominant, followed by the native *B. crenatus*. In the low intertidal zone, *B. crenatus* was the species with the highest abundance. In contrast, the subtidal was dominated by the native *A. improvisus (*Fig. 3a).

Monitoring of barnacle larvae diversity from metabarcoding zooplankton samples at the NIOZ jetty shows that *B. glandula* larvae were already present in the Wadden Sea by November 2023, though the fraction of total reads remained low throughout the monitoring period (Fig 4). *Balanus crenatus* showed the highest fraction of reads throughout this period, followed by *A. modestus. Semibalanus balanoides* mean monthly reads were often lower compared to the other four barnacle species detected in the fieldwork survey. In addition to the barnacles observed during the surveys, metabarcoding also detected larvae o*f Perforatus perforatus*, *Sacculina carcini*, *Verruca stroemia* and *Parasacculina*.

**Fig. 4.**
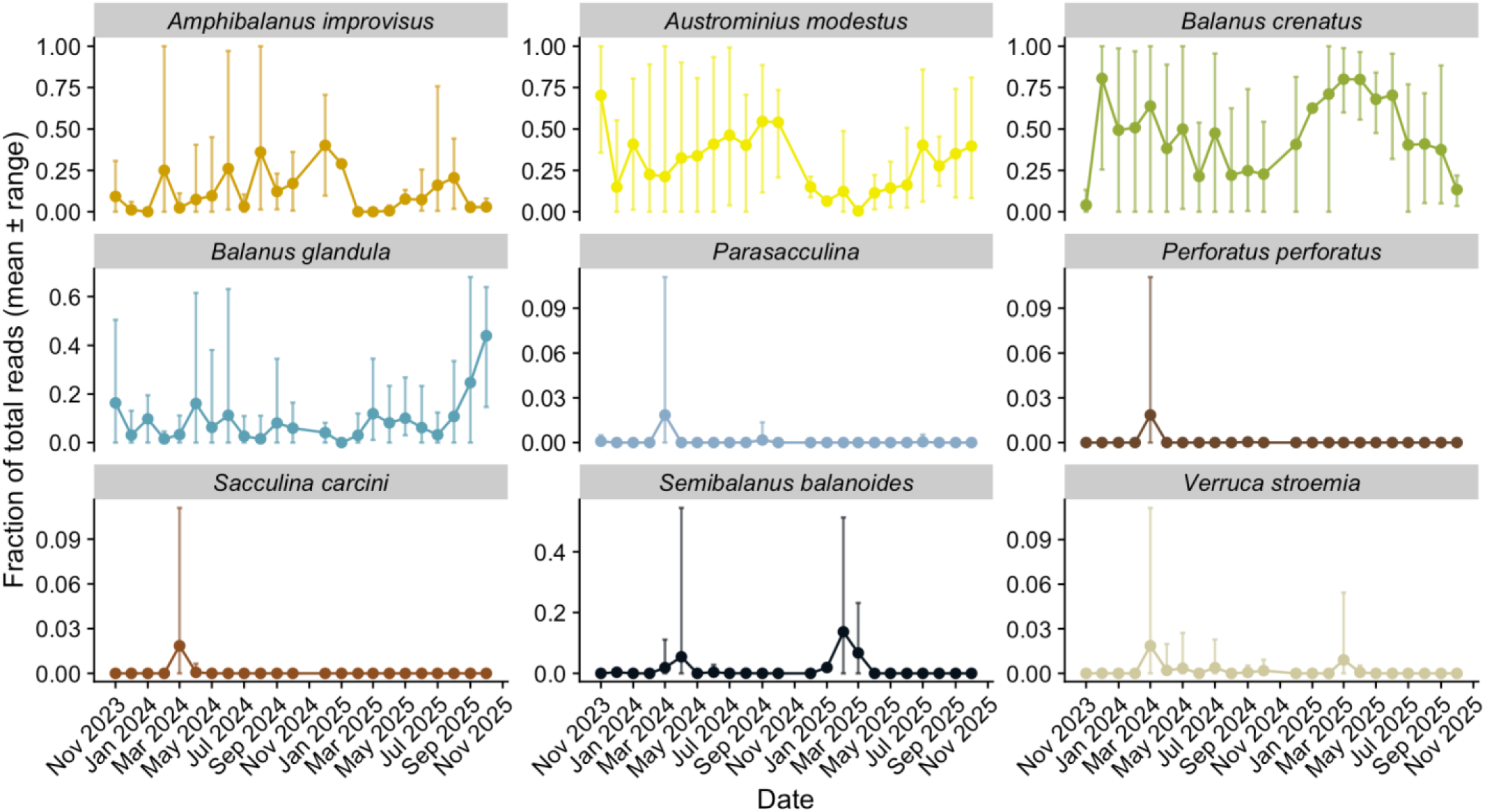
Fraction of total reads (mean ± range) of barnacle species detected through CO1 metabarcoding at NIOZ jetty, Texel (53° 00 ‘06.5”N 4° 47’ 20.5”E) from November 2023 to October 2025. Each bar represents the mean fractional read abundance per species per month. Months with zero reads for a given taxon are not shown.

### 3. Parasite fauna in *Balanus glandula* and other barnacle species

In total, 928 barnacles from 12 sites were dissected to assess parasite infections. Of these, 109 specimens were infected by parasites (overall prevalence of 11.75% across all study sites). Infecting parasites belonged to three different groups: trematodes, cestodes and gregarines (Fig. 5). Trematodes, present as metacercariae (n = 94), had the highest prevalence (overall 10.24%). Eleven barnacles were infected with cestodes (1.18%), and four barnacles harboured gregarine infections (0.43%). Other associated fauna of uncertain or non-parasitic status included nematodes and planarians. Four barnacles were associated with nematodes (0.43%) and two with planarians (0.21%)

**Fig. 5.**
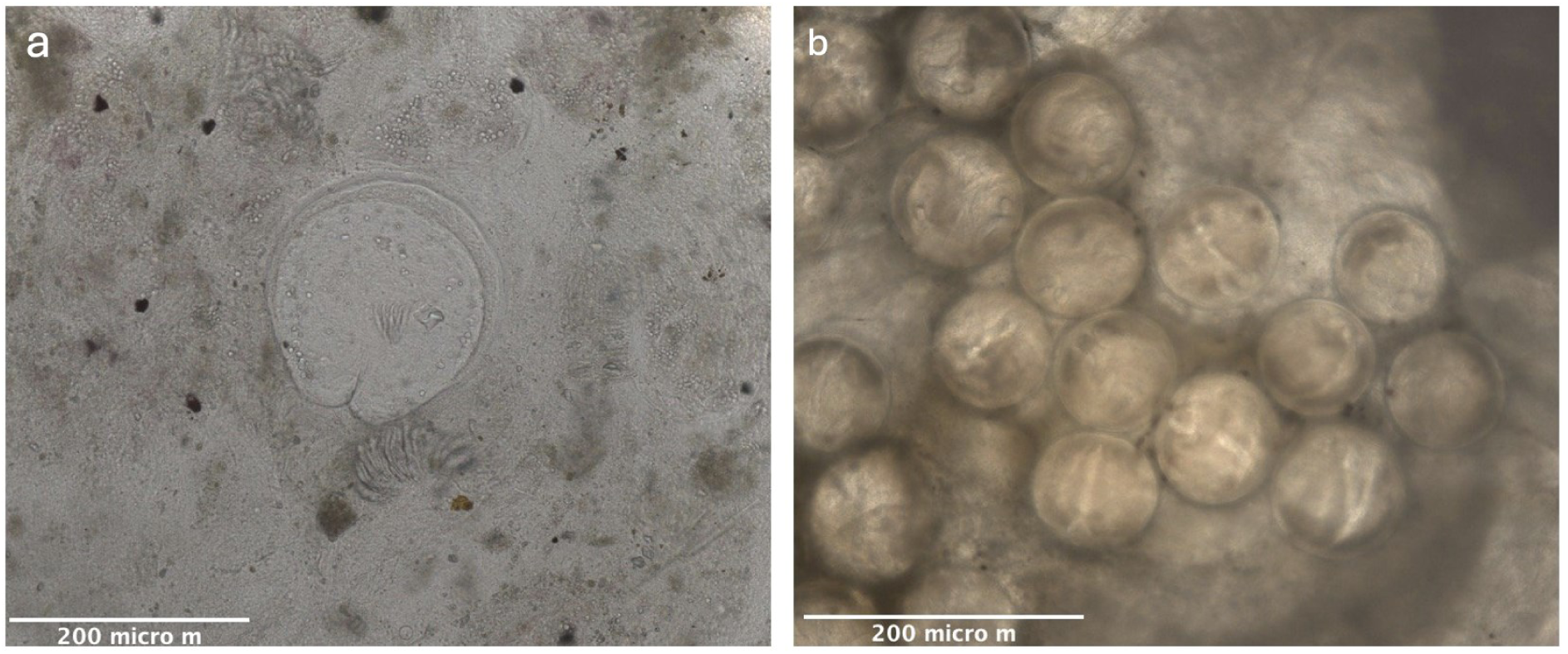
The two main parasite groups found in the barnacles: a) larval cestode cysts and b) encysted larval stages of trematodes

We observed differences in parasite infection prevalences among barnacle species. *Semibalanus balanoides* (n = 180) was only infected by trematode metacercariae at a prevalence of 26.66%. *Balanus crenatus* (n = 115) was infected by metacercariae (2.96%) and by cestodes (0.86%). *Amphibalanus improvisus* (n = 105) was infected by metacercariae (0.95%) and by cestodes (5.71%). *Austrominius modestus* (n = 444) was infected by three types of parasites: metacercariae (9.68%), cestodes (0.90%) and gregarines (0.90%). Gregarine spores were only observed in *A. modestus* from the harbour of List on Sylt*. Balanus glandula* (n = 84) was not found to be infected by any parasite. Nematodes were found in *B. crenatus* (0.86%), *A. modestus* (0.45%) and *B. glandula* (1.18%), although it was unclear whether they were free-living or parasitic. Excluding nematodes of uncertain status, *B. glandula* showed no confirmed parasite infections. Planarians were found in *S. balanoides* (1.11%), but these were free-living (Fig. 6).

**Fig. 6.**
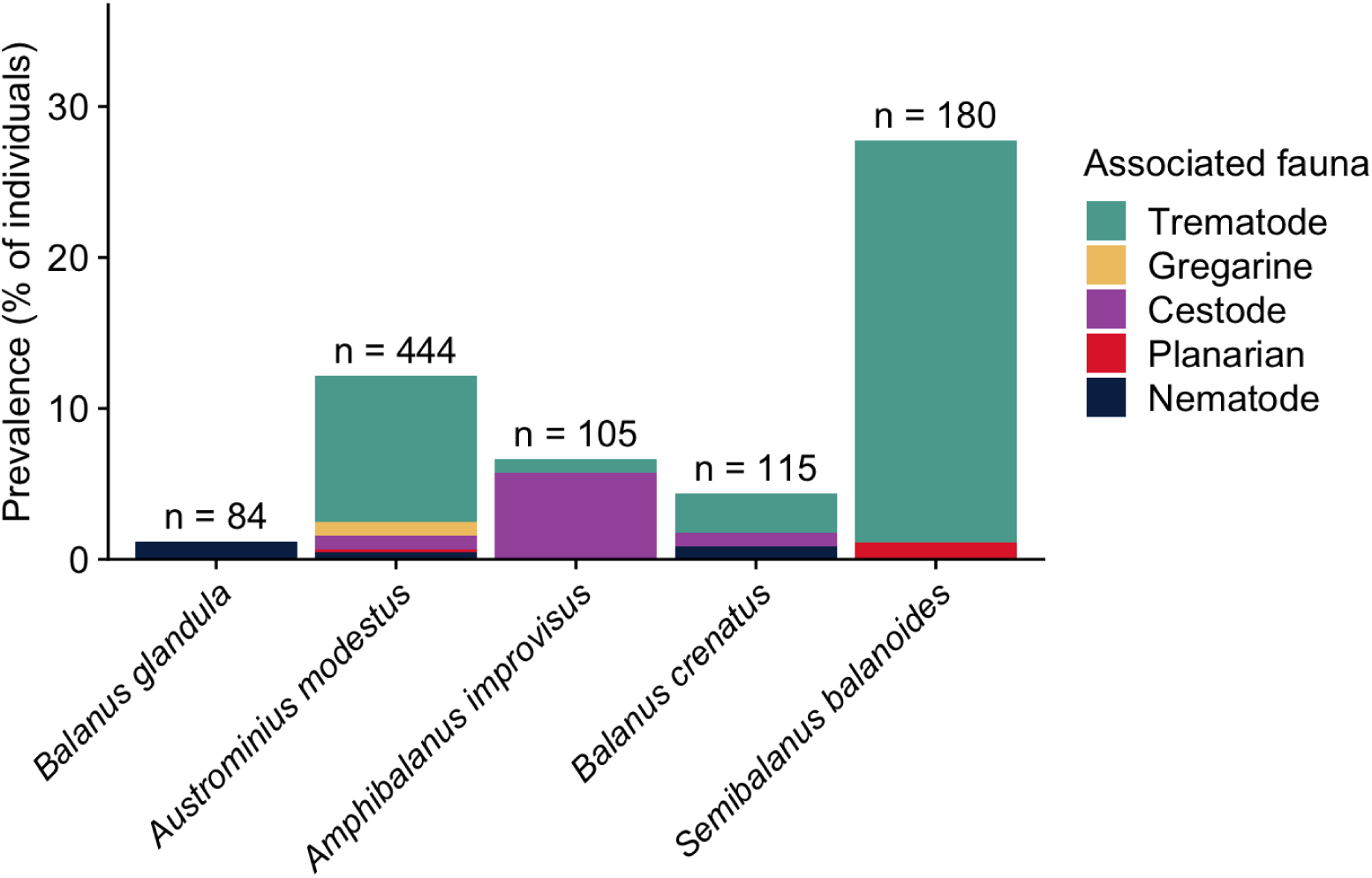
Prevalence of associated fauna observed in the various barnacle species found during field sampling in the Netherlands and Germany.

When comparing infection prevalence of native and invasive barnacles using a binomial regression, native barnacles had a significantly higher likelihood of being infected compared to invasive barnacles (χ²(1) = 5.14, p = 0.023). The null model and the regression model were compared using a log-likelihood ratio test.

Finally, the binomial logistic regression conducted to examine whether the likelihood of infection was related to barnacle species showed overall significant associations (χ²(4) = 57.16, p <0.001). The Tukey post hoc analysis using estimated marginal means showed that *S. balanoides* had a significantly higher probability of infection than *A. modestus* (z = -4.582, df = ∞, p < 0.001). The infection probability differed significantly between *S. balanoides* and *B. crenatus* as well (z = -4.312, df = ∞, p = 0.002). Lastly, *S. balanoides* had a significantly higher probability of infection than *B. glandula* (z = -3.34, df = ∞, p = 0.008). No significant differences were found between the remaining groups.

## Discussion

This study provides the first record of *B. glandula* in the Wadden Sea and extends its known European range northward from the Belgian coast to the Dutch Wadden Sea, with an additional record from a mussel farm on the southern Portuguese coast. *Balanus glandula* was restricted to the high intertidal zone, where *A. modestus* was the dominant species, indicating potential habitat overlap and possible future competition with resident barnacle species. Given its recent introduction to the area, this restricted distribution likely reflects an early stage of colonisation. In contrast to resident species, which harboured trematodes and cestodes, *B. glandula* appeared free of parasites, consistent with parasite release.

### 1. Distribution of *Balanus glandula* in Europe and in the Wadden Sea

Settled individuals of the invasive barnacle *B. glandula* were observed in 2025 in the Dutch Wadden Sea for the first time. The species was initially documented on the European coast in Belgium in 2015 (Kerckhof et al. 2018). It remains unclear when *B. glandula* first established in the Wadden Sea. However, DNA reads matching *B. glandula* have been detected on zooplankton monitoring since November 2023, suggesting that larvae may have been present in the region before the first observation of settled individuals. Potential introduction pathways include hull fouling and ballast water transport, which are well-documented vectors for barnacle introductions (Torres, 2012). In addition, mussel transfers from the Eastern Scheldt to culture sites in the western Wadden Sea represent a plausible regional pathway that warrants further investigation. Cleaning of biofouling from ships happens regularly, but underwater biofouling from harbour structures is often not removed. In three harbours in the Wadden Sea (Den Helder, Groningen and Harlingen), ships large enough to hold ballast water can dock in, but active management of ballast water is lacking (van der Have et al. 2015).

Although *B. glandula* was absent from the sites visited in the German part of the Wadden Sea, we believe that the species has the capacity to spread in this area in the coming decade. Indeed, the closest location in the Netherlands (Lauwersoog) to Germany where *B. glandula* was present is approximately 80 km away from the closest sample point in Germany (Emden), where *B. glandula* was absent. This is in the same order of distance between the island of Texel and Lauwersoog, both with *B. glandula* present, indicating that it is possible that *B. glandula* could be able to cover these distances. The ability of *B. glandula* to disperse is also evident from its spread along the entire Argentine coast in less than 40 years (Schwindt 2007). Additionally, the German Wadden Sea has multiple suitable hard substrates for settlement, including harbour walls, buoys, dams, and other coastal protection structures (Buschbaum et al. 2012; Baptist et al. 2019). Finally, mean sea water temperatures at our northernmost sample points in Sylt (9 °C to 11 °C) fall within *B. glandula’s* wide temperature range (5 °C to 25 °C) (Barnes and Barnes 1956; Schwindt 2007; Nishizaki and Carrington 2014; Rick et al. 2023). Therefore, climatic conditions should not prevent this species from spreading to these latitudes.

Beyond Belgium, we also found records of *B. glandula* from an offshore mussel farm in Southern Portugal in 2018 (Piló et al. 2021). No further range expansion of *B. glandula* on the Portuguese coast has been reported since. The recorded population might have been too small to successfully settle on the Portuguese coast (Piló et al. 2021), or a lack of suitable habitats might be preventing further spread of the species. Alternatively, a lack of screening efforts may explain the apparent absence of range expansion for this species in the area.

### 2. Habitat of *Balanus glandula* and other barnacles in the Wadden Sea

The invasive *B. glandula* was only observed in the high intertidal zone, which matches part of its native range, though it is known to occupy lower intertidal zones as well (Barnes & Barnes, 1953). Its absence from lower intertidal zones may reflect limited detection probability due to the subsampling approach, restricted habitat availability, or biotic interactions with resident barnacle species occupying these zones. However, further studies are needed to determine whether competitive exclusion contributes to the current distribution pattern. Population establishment of *B. glandula* in eastern Hokkaido, Japan, occurred two years after larval recruitment, with peak abundance being reached six years later, suggesting that newly established populations may require considerable time to expand throughout their potential range (Rashidul Alam et al. 2014; Noda and Ohira 2020). Once established, however, *B. glandula* is likely to compete with the dominant invasive *A. modestus* and native *S. balanoides* in the Wadden Sea as it expands, given their overlapping intertidal niches. Its wide thermal tolerance (5–25 °C; Barnes and Barnes 1956; Nishizaki and Carrington 2014) includes the Wadden Sea’s average temperature of approximately 11 °C (Philippart et al. 2024), and in other regions of introduction including Japan, South Africa and Argentina, it has already replaced native and previously invasive species (Schwindt 2007; Geller et al. 2008; Laird and Griffiths 2008). Although ecological impacts can be context-dependent (Kumschick et al. 2015), many interactions are consistent across systems (Buckley and Catford 2016), suggesting *B. glandula* could substantially impact native barnacle communities in the Wadden Sea.

The competitive dominance of the earlier invader, *A. modestus*, is reflected in its widespread colonisation of the high and middle intertidal zones across the Wadden Sea, where it has largely displaced native barnacles (Witte et al. 2010). *B. crenatus*, which was dominant in the low intertidal zone, is known for its limited ability to uptake oxygen from air, compared to high-intertidal species (Foster 1971). The observed distribution of *A. improvisus,* which was only detected in the subtidal, also fitted with its documented habitat preference (Gittenberger et al., 2010). Contrasting with the narrow habitat distribution of the two previous species, *A. modestus* occupies a wide range in the littoral zone and can even inhabit the highest parts (Witte et al., 2010). Although *A*. *modestus* has been observed in subtidal zones before (Watson et al. 2005; O’Riordan et al. 2020), it was absent from the subtidal zone in this study (potentially due to habitat characteristics, as most of the subtidal area consisted of an artificial tree reef in the Wadden Sea). Previously, *A. modestus* was also not found on deeper offshore substrates in the Dutch Wadden Sea (Gittenberger et al. 2009).

Because *A. modestus* inhabits a large portion of the littoral zone, it likely mainly competes with the native barnacle *S. balanoides.* A higher water temperature is considered a key driver of the increasing abundance of *A. modestus* in the Wadden Sea. Indeed, under rising temperatures, larvae of *A. modestus* are less sensitive to food limitation than those of *S. balanoides*, particularly during the cyprid settlement stage (Griffith et al. 2021). On Texel, *A. modestus* was by far the most abundant barnacle in the littoral zone, where it competes with the native S. balanoides for space (Gittenberger et al. 2009). On Sylt, located further north in the Wadden Sea, *A. modestus* has similarly overtaken *S. balanoides* in relative abundance, though at lower proportions, following a period of low abundance that persisted until rising temperatures facilitated its establishment (Witte et al. 2010). This difference is likely due to the slightly colder climate on Sylt, although this could change as heat waves and milder winters become more common in the Wadden Sea (Beukema and Dekker 2020; Philippart et al. 2024).

Not every species picked up by larval metabarcoding is necessarily living there as an established population, as several barnacle species detected were absent from field surveys. *Parasacculina* and *Sacculina carcini* are obligate parasites of decapod crustaceans and would not be expected to occur on hard intertidal substrata (Chan et al. 2021)*. Perforatus perforatus* and *Verruca stroemia* are not characteristic of hard substrata in this region (Gittenberger et al. 2009). Their detection could reflect larval dispersal or transport from adjacent habitats rather than established adult populations. Read abundance derived from metabarcoding data cannot be used as a direct proxy for field abundance. Read counts are influenced by methodological factors including differential primer affinity and DNA degradation. Additionally, species with high larval output relative to their sessile adult abundance, or with larger larval body size, may be disproportionately represented in metabarcoding reads and may also be elevated by species with high larval output relative to their sessile adult abundance.

### 3. Parasite fauna in *Balanus glandula* and other barnacle species

Parasite screening of field-collected barnacles revealed infections by trematodes, cestodes, and gregarines, as well as undetermined association with nematodes and planarians. Additionally, we observed that different barnacle species tend to host different types of parasites, likely reflecting variation in host susceptibility and exposure to infective stages (Arvy and Nigrelli 1970; Vader 1983; Chuang 2021). Trematode metacercarial cysts have been reported in several barnacle species including *A. modestus*, *S. balanoides* and *Chthamalus montagui*, while gregarines have been observed across a broader range of species including *A. improvisus*, *P. perforatus* and *B. glandula* (Priaulx Henry 1938; Colston 2012). At least five types of larval cestodes can infect *S. balanoides* (Chuang 2021; Regel 2024). The isopod *Hemioniscus balani* is widely distributed and has been reported in different barnacles, including, *S. balanoides* and *B. glandula,* on the shores of England, Norway and the United States (Vader 1983; Blower and Roughgarden 1988; Arnott 2001), but was conspicuous by its absence from our samples.

We observed significantly higher prevalence of parasite infections in native barnacles compared to invasive ones, which corroborates the hypothesis that newcomers may be less susceptible to infections, and which may partly explain their success (Goedknegt et al. 2016). None of the dissected invasive *B. glandula* specimens were infected, which can be explained by two potentially complementary mechanisms. First, *B. glandula* presumably lost its parasites when it was introduced in Europe (parasite release; Goedknegt et al. 2016). This possibility is supported by the fact that gregarine infections have been observed in *B. glandula* in its native range (Priaulx Henry 1938), showing that the species is a competent host to some parasites. Second, parasites native to the Wadden Sea have not co-evolved with this newly established host, potentially preventing successful infections (Dunn et al. 2023). No previous-existing studies were found on trematode or cestode infections in *B. glandula*, making it difficult to determine which mechanism is more likely. The lack of data on trematode or cestode infections in *B. glandula*, unfortunately, prevents fully evaluating the possibility of parasite release for this species in the Wadden Sea. Additionally, differences in infection prevalence may not solely be linked with species. Environmental differences among study sites such as tide level and wave exposure, and site-specific variations in the abundance of other intermediate hosts (e.g., snails) may also play roles in shaping parasite transmission and therewith infection levels (Pietrock and Marcogliese 2003). The effects of these additional variables could not be evaluated in our study as not all species were found at every site. Nonetheless, the absence of parasite infections in *B. glandula,* which might reduce host fitness through effects on growth, reproduction and survival, is a competitive advantage over other resident barnacle species, and may facilitate its establishment and spread in coastal areas of western Europe (Goedknegt et al. 2016).

## Conclusions

*Balanus glandula* was recorded in the Wadden Sea for the first time, although its current distribution and abundance remain limited. Given the ability of the species to extend its range after introduction documented in other locations, further spread is expected in the Wadden Sea. The arrival of a second invasive barnacle species in the Wadden Sea (after *A. modestus* several decades ago) might intensify competitive pressure on native barnacle species further, especially on *S. balanoides*. *Balanus glandula* showed no parasite infections in this study, which likely translates to competitive advantages consistent with the enemy release hypothesis. A single observation on the southern Portuguese coast further suggests ongoing range expansion in Europe. Monitoring of *B. glandula* in the Wadden Sea is recommended to track its spread, interactions with native species, and potential parasite acquisition over time.

## Acknowledgements

We would like to thank Robert Twijnstra, Loran Kleine Schaars and Bas de Wit for their technical support at the NIOZ. We are grateful to Dr Annika Cornelius for her assistance during fieldwork on Sylt and for providing additional data from the Rapid Assessment Survey. We also thank Lucìa Irazabal Gonzalez MSc from the Rijksuniversiteit Groningen for making the fieldwork in Lauwersoog possible. In addition, we want to thank Dr Marcel Polling from Wageningen Environmental Research for the metabarcoding analysis of the plankton samples. We want to thank Daisy Greuter for her assistance and support and the reviewers for their comments.

## Author contributions

The study was conceived and designed by Rosalinde van Ooijen in collaboration with David W. Thieltges and Cyril Hammoud. Material preparation, data collection and analysis were performed by Rosalinde van Ooijen, Haiyan He, Annika Cornelius and Cyril Hammoud. The first draft of the manuscript was written by Rosalinde van Ooijen, and all authors commented on previous versions of the manuscript. All authors read and approved the final manuscript.

## Funding

eRAS is funded by the Schleswig-Holstein Wadden Sea National Park Authority and the Lower Saxony Wadden Sea National Park Authority.

## Declarations Conflict of interest

The authors declare no competing financial interests.

## Ethics approval

The authors have no relevant financial or non-financial interests to disclose. The applicable national and institutional guidelines for sampling, care, and experimental use of organisms for the study have been followed.

## Data availability

Data will be made available upon request.

## Notes

### Competing Interest Statement

The authors have declared no competing interest.

