## Supplementary material for "Range expansion of the invasive barnacle *Balanus glandula* into the Wadden Sea, its habitat and parasite fauna compared to established barnacle species": Suppl. Table S2

Table S2. Data for *B. glandula* distribution in Europe. Identification was done based on morphology (M) and/or genetic analysis (GA).

| Country | Region | Location name | Date | Latitude | Longitude | Status | Source | Substrate | Identification |
| --- | --- | --- | --- | --- | --- | --- | --- | --- | --- |
| Belgium | West Flanders | Nieuwpoort | 1/7/2016 | 51,164667 | 2,716667 | absent | Kerckhof et al., 2018 | Bouy | M + GA |
| Belgium | West Flanders | Nieuwpoort | 09/02/2016 | 51,353333 | 2,882000 | absent | Kerckhof et al., 2018 | Bouy | M + GA |
| Belgium | West Flanders | Nieuwpoort | 01/06/2016 | 51,515833 | 2,848667 | absent | Kerckhof et al., 2018 | Bouy | M + GA |
| Belgium | West Flanders | Nieuwpoort | 17/06/2016 | 51,288000 | 2,862167 | absent | Kerckhof et al., 2018 | Bouy | M + GA |
| Belgium | West Flanders | Nieuwpoort | 23/06/2016 | 51,350667 | 2,703667 | absent | Kerckhof et al., 2018 | Bouy | M + GA |
| Belgium | West Flanders | Nieuwpoort | 08/10/2016 | 51,250500 | 2,909500 | present | Kerckhof et al., 2018 | Bouy | M + GA |
| Belgium | West Flanders | Nieuwpoort | 11/10/2016 | 51,417833 | 2,965333 | absent | Kerckhof et al., 2018 | Bouy | M + GA |
| Belgium | West Flanders | Nieuwpoort | 23/10/2016 | 51,399333 | 3,080333 | present | Kerckhof et al., 2018 | Bouy | M + GA |
| Belgium | West Flanders | Nieuwpoort | 23/10/2016 | 51,394500 | 3,080333 | present | Kerckhof et al., 2018 | Bouy | M + GA |
| Belgium | West Flanders | Nieuwpoort | 24/10/2016 | 51,404000 | 3,063333 | present | Kerckhof et al., 2018 | Bouy | M + GA |
| Belgium | West Flanders | Nieuwpoort | 25/10/2016 | 51,248833 | 2,906167 | present | Kerckhof et al., 2018 | Bouy | M + GA |
| Belgium | West Flanders | Nieuwpoort | 16/07/2015 | 51,324000 | 3,113333 | present | Kerckhof et al., 2018 | Bouy | M + GA |
| Belgium | West Flanders | Nieuwpoort | 10/28/2016 | 51,717167 | 2,798500 | absent | Kerckhof et al., 2018 | Bouy | M + GA |
| Belgium | West Flanders | Nieuwpoort | 09/11/2016 | 51,319333 | 3,106167 | present | Kerckhof et al., 2018 | Bouy | M + GA |
| Belgium | West Flanders | Nieuwpoort | 22/11/2016 | 51,221833 | 3,869500 | present | Kerckhof et al., 2018 | Bouy | M + GA |
| Belgium | West Flanders | Nieuwpoort | 22/11/2016 | 51,164667 | 2,716667 | present | Kerckhof et al., 2018 | Bouy | M + GA |
| Belgium | West Flanders | Nieuwpoort | 25/11/2016 | 51,239667 | 2,921500 | present | Kerckhof et al., 2018 | Bouy | M + GA |
| Belgium | West Flanders | Nieuwpoort | 25/11/2016 | 51,240167 | 2,917500 | present | Kerckhof et al., 2018 | Bouy | M + GA |
| Belgium | West Flanders | Nieuwpoort | 25/11/2016 | 51,221000 | 2,872833 | present | Kerckhof et al., 2018 | Bouy | M + GA |
| Belgium | West Flanders | Nieuwpoort | 25/11/2016 | 51,214167 | 2,857333 | present | Kerckhof et al., 2018 | Bouy | M + GA |
| Belgium | West Flanders | Nieuwpoort | 30/11/2016 | 51,389500 | 2,968167 | absent | Kerckhof et al., 2018 | Bouy | M + GA |
| Belgium | West Flanders | Nieuwpoort | 01/12/2016 | 51,289167 | 2,865333 | present | Kerckhof et al., 2018 | Bouy | M + GA |
| Belgium | West Flanders | Nieuwpoort | 09/01/2017 | 51,140333 | 2,573000 | present | Kerckhof et al., 2018 | Bouy | M + GA |
| Belgium | West Flanders | Nieuwpoort | 1/14/2017 | 51,241167 | 2,894167 | present | Kerckhof et al., 2018 | Bouy | M + GA |
| Belgium | West Flanders | Nieuwpoort | 1/18/2017 | 51,301167 | 2,880000 | present | Kerckhof et al., 2018 | Bouy | M + GA |
| Belgium | West Flanders | Nieuwpoort | 1/19/2017 | 51,252833 | 2,862167 | present | Kerckhof et al., 2018 | Bouy | M + GA |
| Belgium | West Flanders | Nieuwpoort | 1/30/2017 | 51,239333 | 2,908333 | present | Kerckhof et al., 2018 | Bouy | M + GA |
| Belgium | West Flanders | Nieuwpoort | 2/2/2017 | 51,270000 | 2,745833 | present | Kerckhof et al., 2018 | Bouy | M + GA |
| Belgium | West Flanders | Nieuwpoort | 2/17/2017 | 51,263667 | 2,893500 | absent | Kerckhof et al., 2018 | Bouy | M + GA |
| Belgium | West Flanders | Nieuwpoort | 2/23/2017 | 51,414500 | 3,184833 | present | Kerckhof et al., 2018 | Bouy | M + GA |
| Belgium | West Flanders | Nieuwpoort | 23/02/2017 | 51,193667 | 2,465333 | absent | Kerckhof et al., 2018 | Bouy | M + GA |
| Belgium | West Flanders | Nieuwpoort | 3/1/2017 | 51,317167 | 3,101500 | present | Kerckhof et al., 2018 | Bouy | M + GA |
| Belgium | West Flanders | Nieuwpoort | 3/20/2017 | 51,374500 | 3,165833 | present | Kerckhof et al., 2018 | Bouy | M + GA |
| Belgium | West Flanders | Nieuwpoort | 4/5/2017 | 51,402333 | 3,303667 | absent | Kerckhof et al., 2018 | Bouy | M + GA |
| Belgium | West Flanders | Nieuwpoort | 5/2/2017 | 51,665667 | 2,878000 | absent | Kerckhof et al., 2018 | Bouy | M + GA |
| Belgium | West Flanders | Nieuwpoort | 5/2/2017 | 51,605000 | 2,805500 | absent | Kerckhof et al., 2018 | Bouy | M + GA |
| Belgium | West Flanders | Nieuwpoort | 5/10/2017 | 51,717167 | 2,798500 | absent | Kerckhof et al., 2018 | Bouy | M + GA |
| Belgium | West Flanders | Nieuwpoort | 5/10/2017 | 51,644333 | 2,753167 | absent | Kerckhof et al., 2018 | Bouy | M + GA |
| Belgium | West Flanders | Nieuwpoort | 6/17/2017 | 51,271667 | 2,979500 | present | Kerckhof et al., 2018 | Bouy | M + GA |
| Belgium | West Flanders | Nieuwpoort | 6/20/2017 | 51,414167 | 3,249667 | present | Kerckhof et al., 2018 | Bouy | M + GA |
| Belgium | West Flanders | Koksijde | 11/28/2017 | 51,118527 | 2,620967 | present | Bouwens 2019 | Groyne | M |
| Belgium | West Flanders | Nieuwpoort | 11/21/2017 | 51,151032 | 2,712921 | present | Bouwens 2019 | Groyne | M |
| Belgium | West Flanders | Westende | 11/14/2017 | 51,17691 | 2,783789 | present | Bouwens 2019 | Groyne | M |
| Belgium | West Flanders | Middelkerke | 11/11/2017 | 51,192821 | 2,819484 | present | Bouwens 2019 | Groyne | M |
| Belgium | West Flanders | Raversijde | 11/27/2017 | 51,207753 | 2,853763 | present | Bouwens 2019 | Groyne | M |
| Belgium | West Flanders | Mariakerke | 11/8/2017 | 51,221431 | 2,884123 | present | Bouwens 2019 | Groyne | M |
| Belgium | West Flanders | Bredene | 11/10/2017 | 51,24899 | 2,95042 | present | Bouwens 2019 | Groyne | M |
| Belgium | West Flanders | Blankenberge | 11/20/2017 | 51,321124 | 3,136017 | present | Bouwens 2019 | Groyne | M |
| Belgium | West Flanders | Duinbergen | 11/23/2017 | 51,348179 | 3,26084 | present | Bouwens 2019 | Groyne | M |
| Germany | Sylt | Ellenbogen | 2025-06-03 | 55,0431543 | 8,4379188 | absent | Own sampling | Oyster bed | M |
| Germany | Sylt | Hörnum coast | 2025-06-05 | 54,7523348 | 8,2792889 | absent | Own sampling | Groyne | M |
| Germany | Sylt | Hörnum harbour | 2025-06-05 | 54,7598151 | 8,295295 | absent | Own sampling | Harbour wall | M |
| Germany | Sylt | List Harbour | 2025-06-04 | 55,0161633 | 8,4394376 | absent | Own sampling | Harbour wall | M |
| Germany | Sylt | Odewatt | 2025-06-02 | 55,0271143 | 8,434686 | absent | Own sampling | Oyster bed | M |
| Germany | Brunsbüttel | Brunsbüttel | 9/5/2025 | 53,88964 | 9,11167 | absent | Neobiota Monitoring program AWI | Harbour walls, stage | M |
| Germany | Brunsbüttel | Brunsbüttel | 9/5/2025 | 53,89605 | 9,14774 | absent | Neobiota Monitoring program AWI | Unknown | M |
| Germany | Brunsbüttel | Brunsbüttel | 9/5/2025 | 53,9116 | 9,17559 | absent | Neobiota Monitoring program AWI | Stones, walls | M |
| Germany | Sylt | List | 9/19/2025 | 55,01615 | 8,44025 | absent | Neobiota Monitoring program AWI | Harbour walls, stage | M |
| Germany | Sylt | List | 9/19/2025 | 55,02226 | 8,441 | absent | Neobiota Monitoring program AWI | Groynes | M |
| Germany | Sylt | List | 9/19/2025 | 54,99393 | 8,38398 | absent | Neobiota Monitoring program AWI | Unknown | M |
| Germany | Dithmarschen | Büsum | 9/2/2025 | 54,12267 | 8,85623 | absent | Neobiota Monitoring program AWI | Groynes | M |
| Germany | Dithmarschen | Büsum | 9/2/2025 | 54,12203 | 8,86504 | absent | Neobiota Monitoring program AWI | Unknown | M |
| Germany | Dithmarschen | Büsum | 9/2/2025 | 54,12332 | 8,86428 | absent | Neobiota Monitoring program AWI | Harbour walls, stage | M |
| Germany | Dithmarschen | Büsum | 9/2/2025 | 54,12565 | 8,86884 | absent | Neobiota Monitoring program AWI | Unknown | M |
| Germany | Sylt | Hörnum | 9/20/2025 | 54,76026 | 8,29608 | absent | Neobiota Monitoring program AWI | Harbour walls, stage | M |
| Germany | Sylt | Hörnum | 9/20/2025 | 54,76078 | 8,29693 | absent | Neobiota Monitoring program AWI | Groynes | M |
| Germany | Sylt | Hörnum | 9/20/2025 | 54,758 | 8,29667 | absent | Neobiota Monitoring program AWI | Unknown | M |
| Germany | Emden | Emden | 9/6/2025 | 53,34324 | 7,19302 | absent | Neobiota Monitoring program AWI | Harbour walls, stage | M |
| Germany | Emden | Emden | 9/6/2025 | 53,33958 | 7,18508 | absent | Neobiota Monitoring program AWI | Harbour walls, stage | M |
| Germany | Emden | Emden | 9/6/2025 | 53,34314 | 7,19315 | absent | Neobiota Monitoring program AWI | Unknown | M |
| Germany | Dithmarschen | Norddeich | 9/7/2025 | 53,62309 | 7,15147 | absent | Neobiota Monitoring program AWI | Harbour walls, stage | M |
| Germany | Dithmarschen | Norddeich | 9/7/2025 | 53,6228 | 7,15473 | absent | Neobiota Monitoring program AWI | Groynes | M |
| Germany | Dithmarschen | Norddeich | 9/7/2025 | 53,62287 | 7,15211 | absent | Neobiota Monitoring program AWI | Unknown | M |
| Germany | Cuxhaven | Cuxhaven | 9/9/2025 | 53,88908 | 8,68425 | absent | Neobiota Monitoring program AWI | Groynes | M |
| Germany | Cuxhaven | Cuxhaven | 9/9/2025 | 53,87518 | 8,70432 | absent | Neobiota Monitoring program AWI | Harbour walls, stage | M |
| Germany | Cuxhaven | Cuxhaven | 9/9/2025 | 53,8891 | 8,68402 | absent | Neobiota Monitoring program AWI | Unknown | M |
| Portugal | Algarve | Lagos | okt-17 | 37,069301 | -8,694594 | present | Piló et al., 2021 | Musselfarm | M |
| Portugal | Algarve | Lagos | apr-18 | 37,069301 | -8,694594 | present | Piló et al., 2021 | Musselfarm | M |
| The Netherlands | Texel | Cocksdorp | 2025-03-26 | 53,1668814 | 4,8748827 | present | Own sampling | Stone | M |
| The Netherlands | Friesland | Lauwersoog | 2025-05-26 | 53,4144526 | 6,2404345 | present | Own sampling | Stone | M |
| The Netherlands | Texel | Mokbaai | 2025-04-14 | 53,0035833 | 4,7779444 | present | Own sampling | Stone | M |
| The Netherlands | Texel | Oudeschild | 2025-03-19 | 53,0480397 | 4,861584 | present | Own sampling | Stone | M |
| The Netherlands | Texel | Paal 15 | 2025-03-03 | 53,0647222 | 4,7202222 | absent | Own sampling | Groyne | M |
| The Netherlands | Texel | Paal 17 | 2025-03-26 | 53,0772764 | 4,7313503 | present | Own sampling | Groyne | M |
| The Netherlands | Vlieland | Waddensea | 2025-4-25 | 53,223 | 5,010306 | absent | Own sampling | Steel | M |
| The Netherlands | Zeeland | Cadzand | 11/22/2017 | 51,382036 | 3,386983 | present | Bouwens 2019 | Groyne | M |
