## Supplementary material for "Range expansion of the invasive barnacle *Balanus glandula* into the Wadden Sea, its habitat and parasite fauna compared to established barnacle species": Suppl. Table S1

Table S1. Field sampling locations. Total amount of barnacles sampled per location, and total amount of barnacles infected per location. Prevalence per barnacle species found at each location. n represents total number of barnacles found (and per site).

|  |  |  |  |  | Prevalence per barnacle species | | | | |
| --- | --- | --- | --- | --- | --- | --- | --- | --- | --- |
| Location | Latitude | Longitude | Total # barnacle sampled | Total # barnacle infected | Amphibalanus improvisus (n = 105) | Balanus crenatus (n = 115) | Semibalanus balanoides (n = 180) | Austrominius modestus (n = 444) | Balanus glandula (n = 84) |
| Cocksdorp | 53.1668814 | 4.8748827 | 43 | 0 | - | 0% (n = 12) | 0% (n = 2) | 0% (n = 8) | 0% (n = 21) |
| Ellenbogen | 55.0431543 | 8.4379188 | 71 | 2 | 12.5% (n = 8) | 2.22% (n = 45) | - | 0% (n = 18) | - |
| Hörnum Coast | 54.7523348 | 8.2792889 | 93 | 60 | - | - | 68.4% (n = 74) | 63.5% (n = 19) | - |
| Hörnum Harbour | 54.7598151 | 8.2952950 | 89 | 0 | - | - | 0% (n = 18) | 0% (n = 71) | - |
| Lauwersoog | 53.4144526 | 6.2404345 | 92 | 1 | 0% (n = 2) | 0% (n = 10) | 2.7% (n = 37) | 0% (n = 38) | 0% (n = 5) |
| List Harbour | 55.0161633 | 8.4394376 | 71 | 7 | - | 5.56% (n = 18) | 0% (n = 4) | 12.2% (n = 49) | - |
| Mokbaai | 53.0035833 | 4.7779444 | 111 | 21 | 0% (n = 1) | 0% (n = 2) | 0% (n = 5) | 22.1% (n = 95) | 0% (n = 8) |
| Odewatt | 55.0271143 | 8.4346860 | 143 | 6 | - | 0% (n = 1) | 0% (n = 33) | 4.58% (n = 109) | - |
| Oudeschild | 53.0480397 | 4.8615840 | 69 | 5 | - | 0% (n = 8) | 0% (n = 7) | 13.51% (n = 37) | 0% (n = 17) |
| Paal 15 | 53.0647222 | 4.7202222 | 4 | 0 | - | 0% (n = 4) | - | - | - |
| Paal 17 | 53.0772764 | 4.7313503 | 47 | 0 | - | 0% (n = 14) | - | - | 0% (n = 33) |
| Vlieland | 53.2230000 | 5.0103060 | 95 | 7 | 6.38% (n = 94) | 0% (n = 1) | - | - | - |
