## Supplementary figures and images for "Range expansion of the invasive barnacle *Balanus glandula* into the Wadden Sea, its habitat and parasite fauna compared to established barnacle species"

### Suppl. Fig S1

**Supplementary Materials**

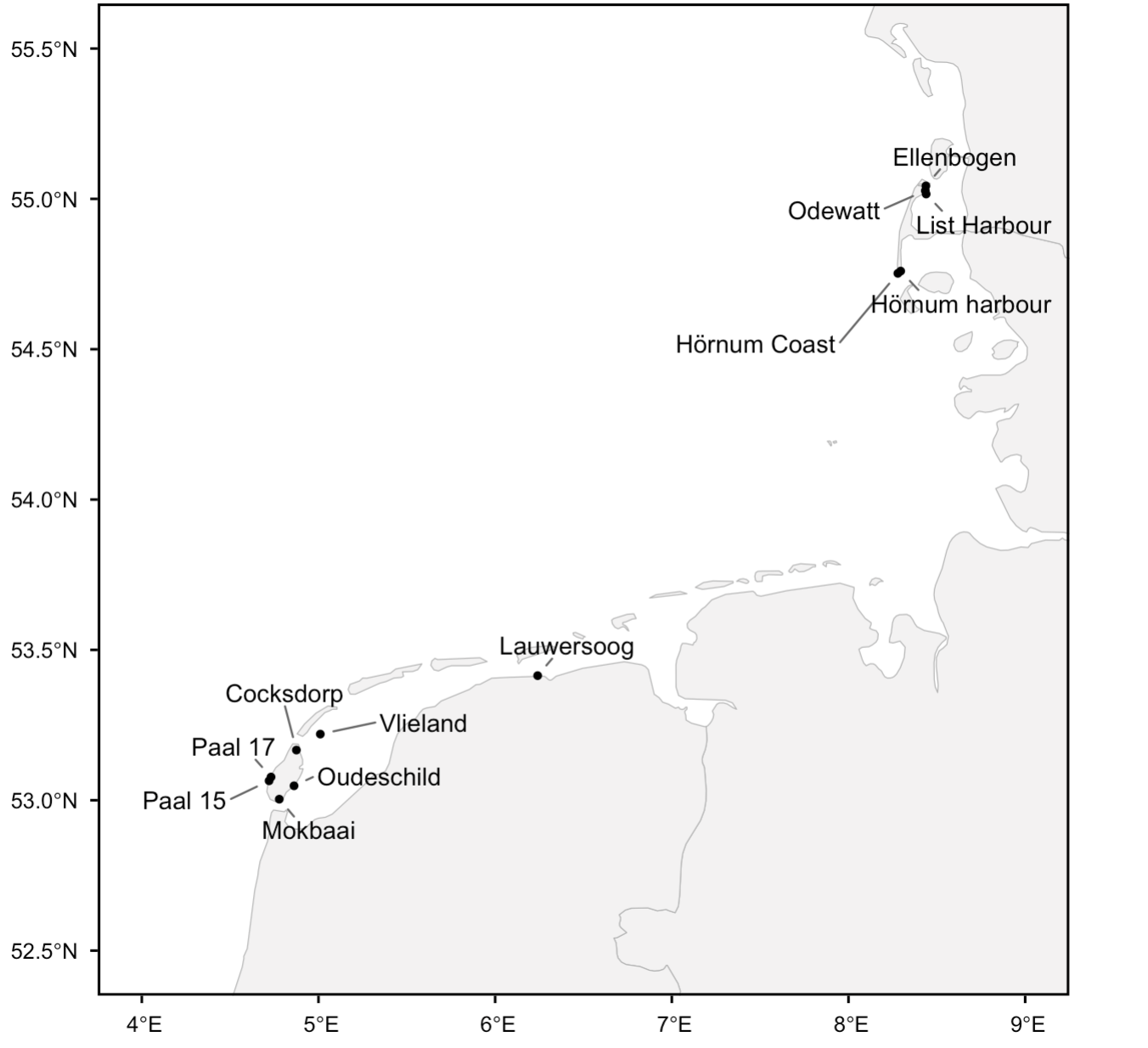

Figure S1. Field sample locations in the Wadden Sea.
